# Cyanobacterial Growth on Municipal Wastewater Requires Low Temperatures

**DOI:** 10.1101/161166

**Authors:** Travis C. Korosh, Andrew Dutcher, Brian F. Pfleger, Katherine D. McMahon

**Author notes:** Corresponding author. 5552 Microbial Science Building, 1550 Linden Drive, Madison, WI 53706, United States.

## Abstract

Side-streams in wastewater treatment plants can serve as concentrated sources of nutrients (i.e. nitrogen and phosphorus) to support the growth of photosynthetic organisms that ultimately serve as feedstock for production of fuels and chemicals. However, other chemical characteristics of these streams may inhibit growth in unanticipated ways. Here, we evaluated the use of liquid recovered from municipal anaerobic digesters via gravity belt filtration as a nutrient source for growing the cyanobacterium *Synechococcus* sp. strain PCC 7002. The gravity belt filtrate (GBF) contained high levels of complex dissolved organic matter (DOM), which seemed to negatively influence cells. We investigated the impact of GBF on physiological parameters such as growth rate, membrane integrity, membrane composition, photosystem composition, and oxygen evolution from photosystem II. At 37°C, we observed an inverse correlation between GBF concentration and membrane integrity. Radical production was also detected upon exposure to GBF at 37°C. However, at 27°C the dose dependent relationship between GBF concentration and lack of membrane integrity was abolished. Immediate resuspension of strains in high doses of GBF showed markedly reduced oxygen evolution rates relative to the control. Together, this suggests that one mechanism responsible for GBF toxicity to *Synechococcus* is the interruption of photosynthetic electron flow and subsequent phenomena. We hypothesize this is likely due to the presence of phenolic compounds within the DOM.

**IMPORTANCE:** Cyanobacteria are viewed as promising platforms to produce fuels and/or high-value chemicals as part of so-called “bio-refineries”. Their integration into wastewater treatment systems is particularly interesting because removal of the nitrogen and phosphorus in many wastewater streams is an expensive but necessary part of wastewater treatment. In this study, we evaluated strategies for cultivating *Synechococcus* strain PCC 7002 on media comprised of two wastewater streams; treated secondary effluent supplemented with the liquid fraction extracted from sludge following anaerobic digestion. This strain is commonly used for metabolic engineering to produce a variety of valuable chemical products and product precursors (e.g. lactate). However, initial attempts to grow PCC 7002 under otherwise standard conditions of light and temperature failed. We thus systematically evaluated alternative cultivation conditions and then used multiple methods to dissect the apparent toxicity of the media under standard cultivation conditions.

## INTRODUCTION

The need to develop non-petroleum based platforms for fuel and chemical production is driving researchers to explore alternatives that harness renewable energy sources while minimizing other environmental impacts such as freshwater depletion, eutrophication, and the use of arable land for non-food production. Cyanobacteria are particularly attractive such platforms due to their genetic tractability, rapid growth rates, halotolerance, and ability to be grown on non-productive land with simple nutrient requirements (1, 2). According to published life cycle assessments, a large portion of the associated costs of culturing photoautotrophs are tied to upstream costs, such as CO_2_ delivery and fertilizer application (3). High phosphorus/nitrogen removal rates and energy efficiencies have been reported for photobioreactor and open pond cultivation systems using wastewater streams rich in nitrogen and phosphorus (4, 5). Therefore, it may be possible to offset the requirement for fertilizer by reclaiming nutrients from wastewater. This approach could yield the sought-after non-petroleum based alternative while also providing a more effective means of nutrient and metal removal than conventional wastewater treatment (6–8). Side-streams from common wastewater treatment facilities such as supernatants or filtrates from solids-separation processes are particularly promising nutrient sources, assuming that cyanobacterial strains can efficiently use them.

Of the many streams available in common wastewater facilities, the liquid fraction of anaerobic digestate is thought to be the most attractive nutrient source (8–11). Although digestate is rich in the inorganic constituents necessary for growth, it also contains dissolved organic matter (DOM) that has been shown to limit photosynthetic activity (12). DOM is a heterogenous mixture of aliphatic and aromatic compounds derived from the decomposition of living organisms (13). The chemical nature of wastewater-derived DOM is largely governed by the type of treated waste and the treatment process, but it is largely comprised of hydrophilic, fulvic, and humic substances (14, 15). Humics can induce damaging permeability in model and bacterial membranes (16, 17). Various studies have also demonstrated that fulvic and humic acids can enhance the solubility of many organic compounds (18), which in turn would augment their bioavailability and potential membrane permeability. Many of these compounds are also photo-reactive, producing toxic hydrogen peroxide and hydroxyl radicals (19, 20), which is of significant concern for phototroph cultivation.

The mode of DOM toxicity is thought to involve interactions with the protein-pigment complex of photosystem II (PSII) in photosynthetic organisms, although the exact molecular mechanism remains unclear (21). When the rate of light induced damage to PSII exceeds its rate of repair, growth suppression and chlorosis result from the phenomena known as photoinhibition (22). When damage by photoinhibition is sufficient to hamper the natural ability to consume electrons generated by photosynthesis, reactive oxygen species (ROS) are concomitantly produced as an undesired by-product. Prolonged oxidative stress halts protein translation through oxidation of specific cysteine residues in the ribosomal elongation factors (23). Given these findings, it is increasingly evident that under conditions of sustained stress, regulation of electron flow is critical to maintain homeostasis in photosynthetic organisms (24, 25). Thus, it is important to understand the mechanisms by which DOM may be interrupting electron flow in order to capitalize on the potential of cyanobacteria to remediate wastewater and generate high-value chemicals.

In this study, we tested the practicality of using combined streams from a municipal wastewater plant as a nutrient source for cyanobacterial cultivation. We used the euryhaline cyanobacterium *Synechococcus* sp. strain PCC 7002 (PCC 7002) due to its exceptional tolerance to high light intensity, salt, and other environmental stresses (26, 27). Initial attempts to grow PCC 7002 in this nutrient source under standard environmental conditions for this strain (1% (v/v) CO_2_, a temperature of 37°C, and illumination of 200 µmol photons m^−2^s^−1^) resulted in photobleached (white-yellow) cultures. In an effort to explain this observation while developing more feasible cultivation conditions, we assessed the effects of many environmental parameters during PCC 7002 cultivation in wastewater-based media by monitoring changes in growth rate, photopigment abundance, oxygen evolution rates, membrane integrity, and membrane composition. High GBF concentrations were associated with elevated DOM levels. Decreased photosynthetic oxygen production rates were also noted upon exposure to high levels of GBF. We observed marked membrane permeability, photosystem degradation, low growth rates, and ROS production in cultures exposed to GBF at 37°C. In comparison, at 27°C the cultures grown on GBF had reduced membrane permeability, robust growth rates, as well as high levels of total fatty acids and an elevated unsaturated fatty acid content relative to control. This suggests bioavailability of the photoinhibitory compound in GBF is governed by changes in membrane content and composition that occur during growth.

## MATERIALS AND METHODS

### Media and Growth Conditions

*Synechococcus* sp. strain PCC 7002 was obtained from the Pasteur Culture Collection of Cyanobacteria. Experiments were performed with a strain of PCC 7002 harboring the *aaC1* gene encoding a gentamicin resistance marker in the A2842 locus (*glpK*) to maintain axenic cultures. Strains were grown and maintained on solid media composed of Medium A^+^ (28) supplemented with 5 µM NiSO_4_ (29) with 1.5% Bacto-Agar. Strains were cultured from a single colony in 250 ml baffled flasks with 50 mL media with 1% CO_2_-enriched air at 150 rpm in a Kuhner ISF1-X orbital shaker. Temperature was maintained at 37°C or 27°C and light intensity was fixed at approximately 200 µmol photons m^−2^s^−1^ or 100 µmol photons m^−2^s^−1^ via a custom LED panel. Strains were pre-acclimated to the culture conditions overnight before inoculating in fresh media. Optical density at 730 nm was measured in a Tecan M1000 plate reader.

Wastewater-derived media was obtained from the Nine Springs Wastewater Treatment Plant (Dane County, Wisconsin, USA). The plant is configured for biological nutrient removal via a modified University of Cape Town process with no internal nitrate recycling and stable performance yielding high secondary effluent quality (total phosphorus < 1 mg P/L, ammonia < 1 mg N/L, nitrate ~ 15 mg N/L) (30). Anaerobic digesters are used for solids stabilization and the resulting digested material is passed over a gravity belt filter for dewatering. The system includes an Ostara WASSTRIP process to recover phosphorus. This filtrate (GBF) served as the primary source of phosphorus and reduced nitrogen for our cultures, and effluent from the post-mainstream secondary treatment clarifier (secondary effluent) served as a diluent. GBF was gravity filtered through a paper coffee filter to remove any exceptionally large flocs, then stored at −80°C until use. Secondary effluent was collected one to four days before each experiment and held under refrigeration at approximately 2°C. Experimental media was comprised primarily of secondary clarifier effluent and GBF, combined in different proportions. Unless otherwise noted, GBF was used at a concentration of 12.5% (v/v) in secondary effluent, supplemented with trace metals and vitamin B12 at the concentrations found in Medium A^+^, as well as KH_2_PO_4_ at a molar ratio of 1:32 soluble reactive phosphorus (SRP) to bioavailable nitrogen (the sum of NH_4_^+^ and NO_3_^−^). All media was buffered with Tris-HCl and adjusted to pH 8.0 with potassium hydroxide or hydrochloric acid before sterilization by autoclaving, and gentamycin was added at working concentrations (30 µg/mL) after cooling.

### Staining, Flow Cytometry, and Fluorescence Measurements

The membrane permeability of cyanobacteria cells was recorded by a flow cytometer (FACSCalibur, BD Biosciences, San Jose, CA, USA). After growth in the respective media, cells were centrifuged (2 minutes, 5000 RCF), decanted, and resuspended in 1 mL of Tris-Buffered Saline (TBS) solution (pH 8.0). To identify membrane-compromised cells, SYTOX Green (Life Technologies) was also added to each sample (1 µM). SYTO 59 (Life Technologies) was added to each sample (1 µM) as a nucleic acid counterstain. As a control for a permeabilized membrane, cells were resuspended in 190 proof ethanol. SYTOX green fluorescence was visualized using 488 nm laser excitation and emission area was read using a 530/30 nm bandpass filter. The 633 nm laser coupled with a 661/16 bandpass filter was used for SYTO 59 visualization. Analysis of the cytometric data was carried out with CellQuest Pro (BDBiosciences, San Jose,CA, USA) software.

To assess reactive oxygen species production, cells (OD_730_ = 1) were incubated overnight in Medium A^+^, 12.5 % (v/v) GBF, 12.5 % (v/v) GBF + 1 mM Dithiothreitol (DTT), 12.5 % (v/v) GBF + 1 mM N-acetylcysteine (NAC), or Medium A^+^ + 100 µM methyl viologen as a positive control at 37°C with 5% CO_2_ at a light intensity of 200 µmol photons m^−2^s^−1^. Cells were washed in TBS, and either Sytox Green (1 µM) or CellROX Orange reagent (Life Technologies) (5 µM) was added. After 30 min incubation in darkness at 37°C, fluorescence was measured (Ex/Em 545/565 nm for Cell ROX Orange) and (Ex/Em 504/523 nm for Sytox Green) in a Tecan M1000 plate reader.

### GBF Characterization

SRP, ammonia, nitrate, and nitrite concentrations were determined for all secondary clarifier effluent and GBF samples used in these experiments. In addition, the GBF was tested for total suspended solids (TSS), volatile suspended solids (VSS) total solids (TS), and chemical oxygen demand (COD). Ammonia, SRP, and COD concentrations were determined by colorimetric tests using reagents from Hach. Nitrate and nitrite were determined using high performance liquid chromatography (Shimadzu) with a C18 column and photodiode array detector (31). TSS, VSS, and TS were measured according to Standard Method 2540 D, 2540 E, and 2540 B, respectively, with 47mm diameter glass fiber filters (Whatman) used for TSS and VSS (32). Fluorescence EEM measurements and UV-Vis absorbance scans were conducted using a M1000 Tecan plate reader using a UV-transparent plate (Costar 3635). Fluorescence intensity was normalized to quinine sulfate units (QSU), where 1 QSU is the maximum fluorescence intensity of 1 ppm of quinine sulfate in 0.1 N H_2_SO_4_ at Ex/Em = 350/450. Rayleigh scatter effects were removed from the data set.

### Biochemical analyses

Batch cultures were further assayed for dry cell weight (DCW), fatty acid content, oxygen evolution rates, and chlorophyll a content. Cells were concentrated by centrifugation, washed in TBS, and lyophilized overnight to obtain dry cell weights (DCW). Fatty acids from approximately 10 mg of DCW with 10 mg ml^−1^ pentadecanoic acid as an internal standard were converted to methyl-esters, extracted with n-hexane and analyzed by GC-FID on a Restek Stabilwax column (60m, 0.53 mm ID, 0.50 µm) (33). Photosynthetic oxygen evolution from whole cells was measured with a Unisense MicroOptode oxygen electrode with 10 mM NaHCO_3_ illuminated with a slide projector at photosynthetic photon flux densities ranging from 76–2700 µmol photons photons m^−2^ s^−1^ for 30 min at room temperature (34). Cells were collected by centrifugation and resuspended in the appropriate media to give an OD_730_ = 1.0. Chlorophyll a measurements were done via a 100% chilled methanol extraction procedure (35). Chlorophyll a was calculated via the following equation: Chl_a_ = 16.29 * A^665^ - 8.54 * A^652^ (36).

## RESULTS

### GBF and secondary effluent characteristics

We measured nutrient concentrations from the batches of GBF collected over the 6 month experimental period (Table 1 and Table 2), to ascertain if it was a stable and reliable nutrient source for cultivating the cyanobacteria. Sampling points from the Nine Springs Wastewater Treatment Plant (Dane County, Wisconsin, USA) are shown in Figure 1. In order to calculate nutrient stoichiometries, we took the sum of NH_3_-N and NO_3_-N to be the bioavailable N. Nutrient levels in the GBF were markedly more variable than in the secondary effluent. The average molar ratio of bioavailable N to SRP was 35 ± 7 in GBF (12.5% v/v) diluted with secondary effluent, as compared to 32 in Medium A^+^.

**Table 1.** Nutrient composition of batch of 100% GBF used for subsequent experiments

| Analyte<sup>a</sup> | Concentration<sup>b</sup> (mg L<sup>-1</sup>) (±SD) |
| --- | --- |
| NH<sub>3</sub>-N | 1180 ± 135 |
| NO<sub>3</sub>-N | 7.5 ± 0.04 |
| SRP | 78 ± 10 |
| COD | 735 ± 4 |
| TSS | 467 ± 123 |
| VSS | 18 ± 2 |
| TS | 2350 ± 36 |
| TS (Glass Fiber Filtered) | 2030 ± 42 |
| TS (0.45 um membrane filtered) | 2000 ± 151 |

<sup>a</sup> SRP-soluble reactive phosphorus, COD-chemical oxygen demand, TSS- total suspended solids, VSS – volatile suspended solids, TS-total solids
<sup>b</sup> Values shown represent the mean ± standard deviation (SD) of at least three technical replicates.

**Table 2.** Characteristics of GBF and secondary effluent over the 6 month experimental period.

| Source | Analyte | Concentration<sup>a</sup> (mg L<sup>-1</sup>) (±COV) |
| --- | --- | --- |
| GBF | NH<sub>3</sub>-N | 920 ± 22% |
|  | NO<sub>3</sub>-N | 3.9 ± 43% |
|  | SRP | 54 ± 27% |
| Secondary Effluent | NH<sub>3</sub>-N | n.d. |
|  | NO<sub>3</sub>-N | 19.3 ± 6.6% |
|  | SRP | n.d. |

<sup>a</sup> Values shown represent the mean ± coefficient of variation (COV) of at least three technical replicates.

**Figure 1.**
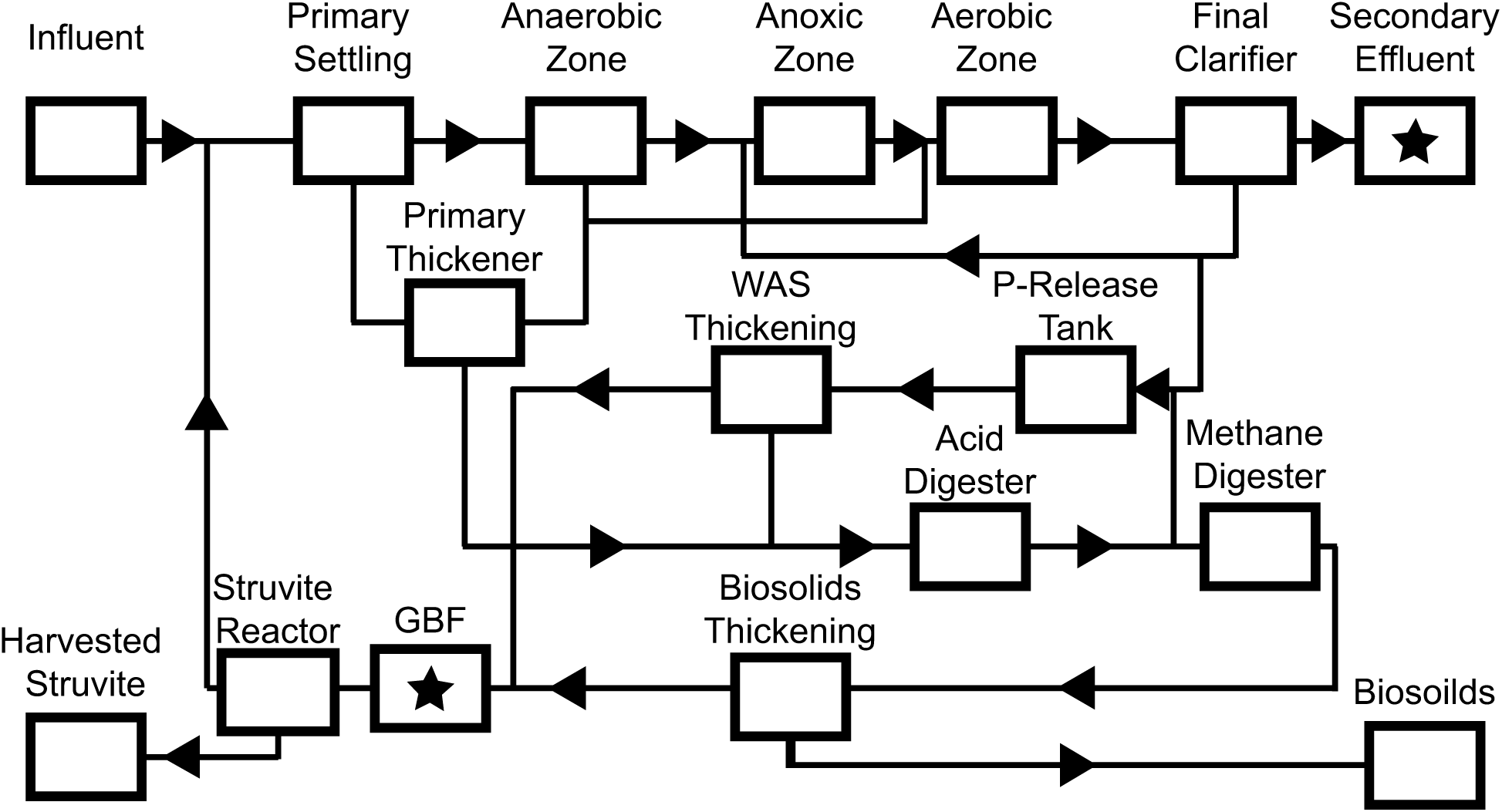
Flow diagram and nutrient streams obtained from the Nine Springs Wastewater Treatment Plant (Dane County, Wisconsin, USA). Stars indicate sampling points.

We also measured DOM quality in the secondary effluent and GBF using excitation−emission matrix (EEM) fluorescence spectroscopy (37) because we hypothesized that DOM was linked to toxicity during cultivation, as has been shown in prior studies (16, 17, 38). EEM fluorescence spectroscopy revealed that secondary effluent contained diffuse constituents, including humic [excitation wavelengths (>280 nm) and emission wavelengths (>380 nm)] and fulvic acid-like [excitation wavelengths (<250 nm) and emission wavelengths (>350 nm)] spectra relative to Medium A^+^ (Figure 2). The distinction between the two substances is historically based on solubility (39), but compositionally, fulvic acids contain more acidic functional groups than humic acids (40). The fulvic acid content in media preparations rose with increasing GBF concentrations. Additionally, at high concentrations of GBF, we detected absorbance in a distinct region [excitation wavelengths (270–290 nm) and emission wavelengths (340–400 nm)] that points to the presence of soluble microbial products, which include aromatic amino acids, carbohydrates, or phenols (37, 41). Prior work has shown that the GBF stream of the Nine Springs Treatment Plant contained high molecular weight DOM, which was dissimilar in composition to other internal streams (42)

**Figure 2.**
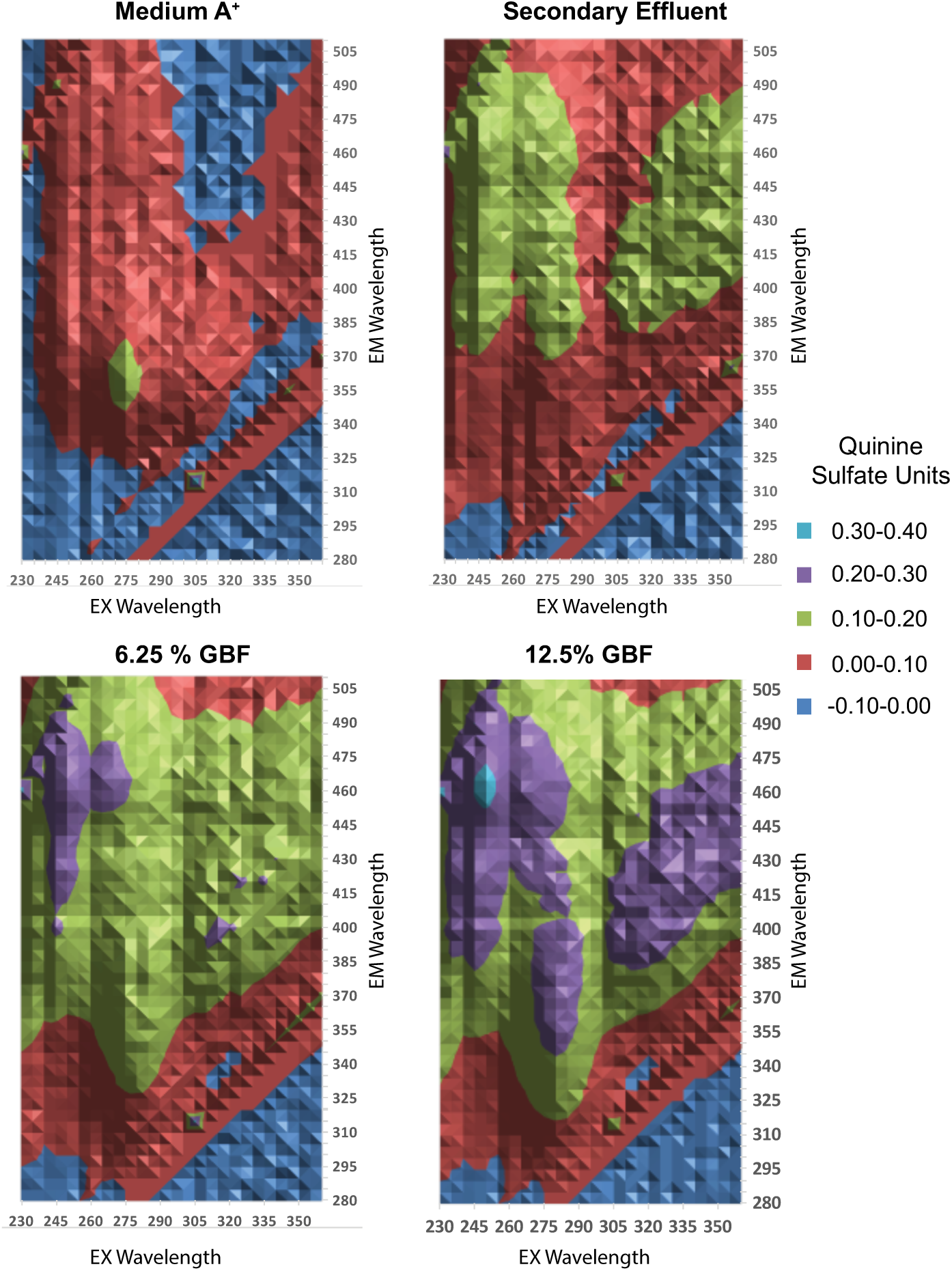
Excitation Emission Matrix of (A) Medium A+, (B) Secondary Effluent, (C) 6.25% GBF, and (D) 12.5% GBF. Fluorescence values were normalized to 1 ppm of quinine sulfate in 0.1 N H_2_SO_4_ at Ex/Em = 350/450 nm.

### Dose-dependent tolerance to GBF is a function of growth temperature

To evaluate the effects of GBF dosage on PCC 7002 physiology, we measured biomass accumulation (Figure 3) and growth rates (Table 3) in media with GBF concentrations ranging from 6.25%–12.5% (v/v), as a function of temperature (27°C vs 37°C) and light intensity (100 µmol photons m^−2^ s^−1^ vs 200 µmol photons m^−2^ s^−1^). Medium A^+^ served as a control. Higher temperatures depressed growth rates in GBF-based media. At 37°C, growth rates with 6.25% GBF at both 100 µmol photons m^−2^ s^−1^and 200 µmol photons m^−2^ s^−1^ were most comparable to Medium A^+^. Higher GBF concentrations had a more extreme effect on growth rates. However, this dose-dependent effect of GBF on growth rate was abolished when the cultivation temperature was lowered to 27°C. At 200 µmol photons m^−2^ s^−1^ and 27°C, GBF cultures grew twice as slowly as the control and there was no significant difference between the tested GBF concentrations. Under 100 µmol photons m^−2^ s^−1^and 27°C, growth rates were comparable across media conditions. Thus, successful cultivation using the more concentrated GBF media required adjusting both the light and temperature regimes.

**Table 3.** Growth Rates with varying Temperature and Light Intensity

| Media | Light Intensity<br>($\mu\text{mol photons m}^{-2} \text{ s}^{-1}$) | Temperature<br>($^{\circ}\text{C}$) | Growth Rate ($\pm\text{SD}$)<br>($\text{OD day}^{-1}$) |
| --- | --- | --- | --- |
| A<sup>+</sup> | 200 | 37 | $0.66 \pm 0.04$ |
| 6.25% GBF | 200 | 37 | $0.58 \pm 0.05$ |
| 12.5% GBF | 200 | 37 | $0.05 \pm 0.01$ |
| A<sup>+</sup> | 100 | 37 | $0.28 \pm 0.01$ |
| 6.25% GBF | 100 | 37 | $0.24 \pm 0.01$ |
| 9.9% GBF | 100 | 37 | $0.11 \pm 0.02$ |
| 12.5% GBF | 100 | 37 | $0.02 \pm 0.00$ |
| A<sup>+</sup> | 200 | 27 | $0.73 \pm 0.08$ |
| 6.25% GBF | 200 | 27 | $0.32 \pm 0.03$ |
| 9.9% GBF | 200 | 27 | $0.42 \pm 0.06$ |
| 12.5% GBF | 200 | 27 | $0.28 \pm 0.04$ |
| A<sup>+</sup> | 100 | 27 | $0.25 \pm 0.01$ |
| 6.25% GBF | 100 | 27 | $0.26 \pm 0.02$ |
| 9.9% GBF | 100 | 27 | $0.24 \pm 0.01$ |
| 12.5% GBF | 100 | 27 | $0.26 \pm 0.02$ |

<sup>a</sup> Linear growth rates of cultures grown in Medium A<sup>+</sup> or GBF with the appropriate treatment under continuous illumination ( $200 \mu\text{mol photons m}^{-2} \text{ s}^{-1}$ or $100 \mu\text{mol photons m}^{-2} \text{ s}^{-1}$ ) with 1 % CO<sub>2</sub> at 37°C or 27°C. The values represent the mean ± SD of biological triplicates.

**Figure 3.**
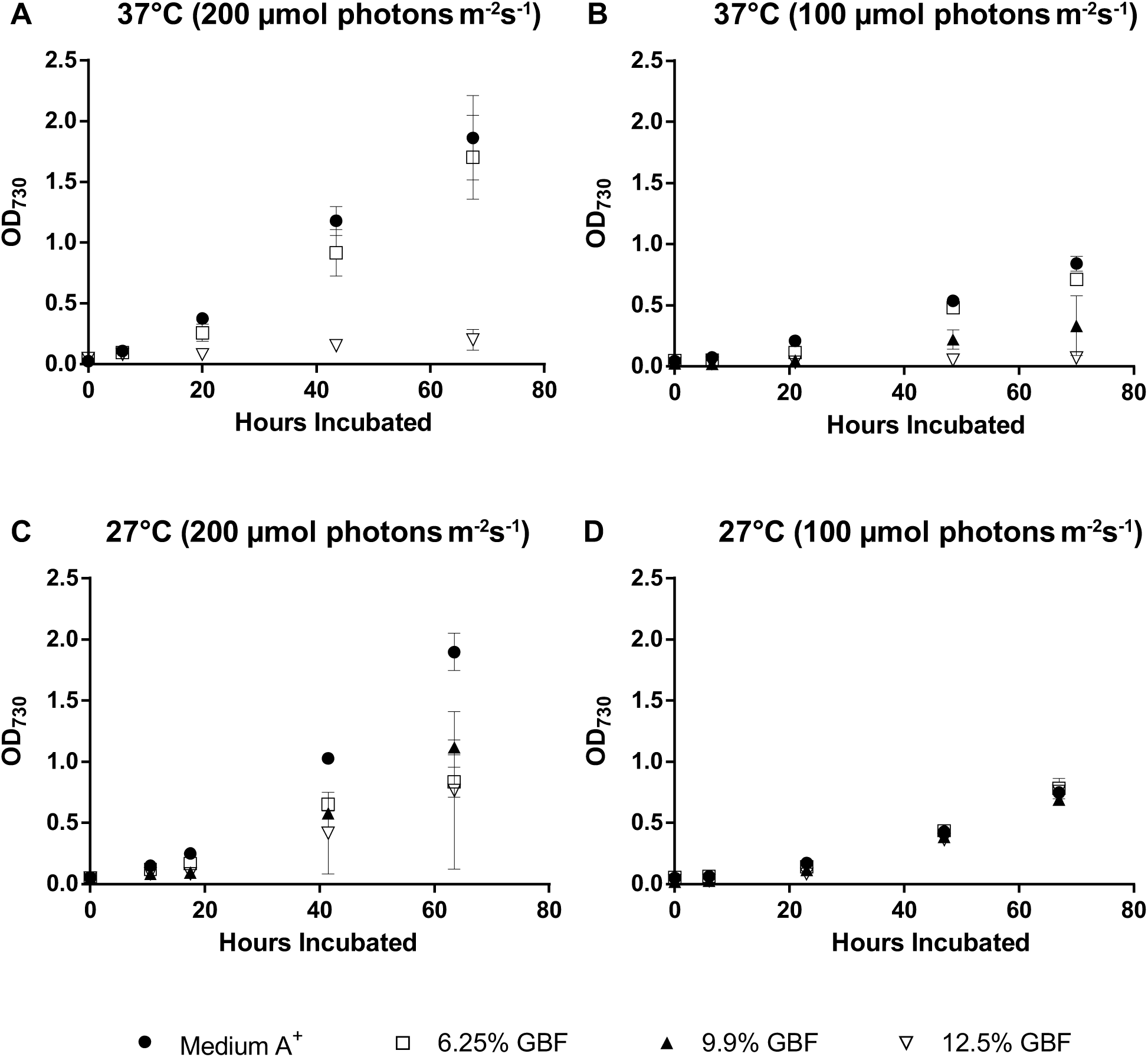
Biomass accumulation for cultures grown in Medium A^+^, 6.25 %, 9.9 %, or 12.5 % (v/v) GBF media with 1 % CO_2_ at (A) 37°C and 200 µmol photons m^−2^ s^−1^, (B) 37°C and 100 µmol photons m^−2^ s^−1^, (C) 27°C and 200 µmol photons m^−2^ s^−1^, or (D) 27°C and 100 µmol photons m^−2^ s^−^1. The values represent the mean ± SD of biological triplicates.

### Dose-dependent membrane permeability of GBF is a function of growth temperature

We wondered whether the decreased growth rates in GBF at 37°C was a result of membrane permeability, given the known effect of humic acids on membrane integrity (16, 17). To track the dynamics of GBF induced membrane permeability, we employed forward scatter flow cytometry using SYTO 59 as a counterstain to identify cells, which were subsequently visualized for membrane permeability using Sytox Green. As Sytox Green is a membrane impermeable dye that fluoresces when binding to nucleic acids (43), fluorescence would indicate compromised outer membrane structures. Two distinct phases were identified upon exposure to GBF, which we interpreted as initial and chronic membrane permeability (Figure 4). Initial membrane permeability was defined as the Sytox Green positive events for samples analyzed within the first 10 hours of growth, while chronic membrane permeability accounted for the Sytox Green positive events during subsequent time points. As was the case for growth rate, a relationship between both initial and chronic membrane permeability and increasing GBF concentrations was found at 37°C (Figure 4a). While we still detected considerable initial membrane permeability with GBF exposure at 27°C, chronic membrane permeability decreased over time, mostly likely to the increase in biomass (Figure 3). Altogether, this suggested that there was a temperature dependent adaptation that ameliorated the susceptibility of cultures to GBF induced membrane permeability.

**Figure 4.**
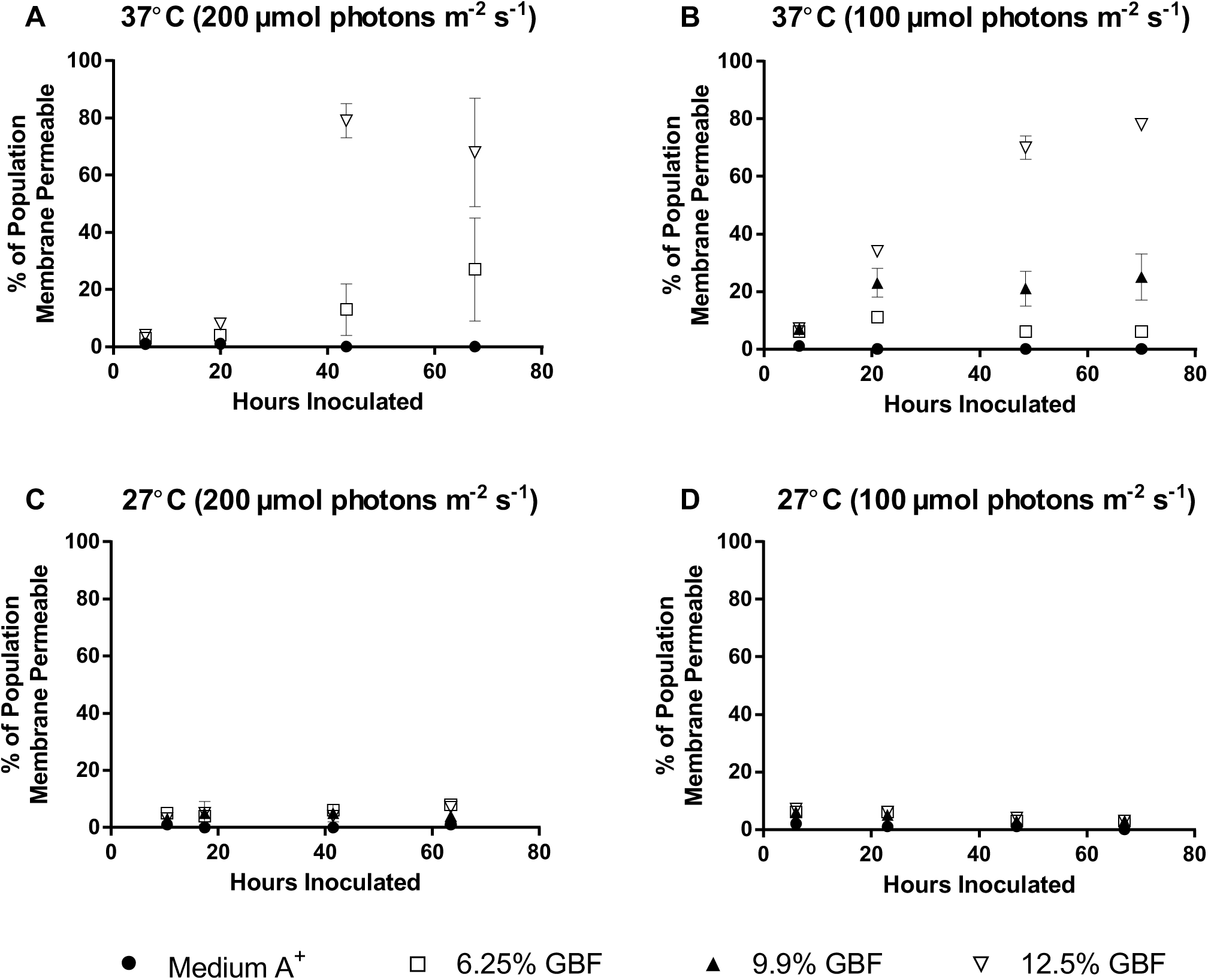
Membrane permeability of cultures grown in Medium A^+^, 6.25 %, 9.9 %, or 12.5 % (v/v) GBF media with 1 % CO_2_ at (A) 37°C and 200 µmol photons m^−2^ s^−1^, (B) 37°C and 100 µmol photons m^−2^ s^−1^, (C) 27°C and 200 µmol photons m^−2^ s^−1^, or (D) 27°C and 100 µmol photons m^−2^ s^−^1. The values represent the mean ± SD of biological triplicates.

### Exposure to GBF at high temperatures generates radicals and destroys the photosynthetic pigments

We directly measured the ROS content and membrane permeability in response to overnight GBF exposure at 37°C using the fluorophores Sytox Green and CellROX Orange (Figure 5). We examined the capacity of reducing agents Dithiothreitol (DTT) or N-acetylcysteine (NAC) to quench the media toxicity, since they have anti-oxidant properties due to the direct reduction of disulfide bonds or as precursors for the anti-oxidant glutathione (44). To measure their effect on the ROS production and culture viability after GBF exposure, we assessed ROS content and membrane integrity with either concurrent addition or preincubation of these quenching compounds in the diluted (12.5% v/v) GBF media. Addition of 100 µM methyl viologen to Medium A^+^ served as a positive control. Exposure of cells to 12.5%-GBF media resulted in marked ROS production and membrane permeability, which no thiol treatment alleviated.

**Figure 5.**
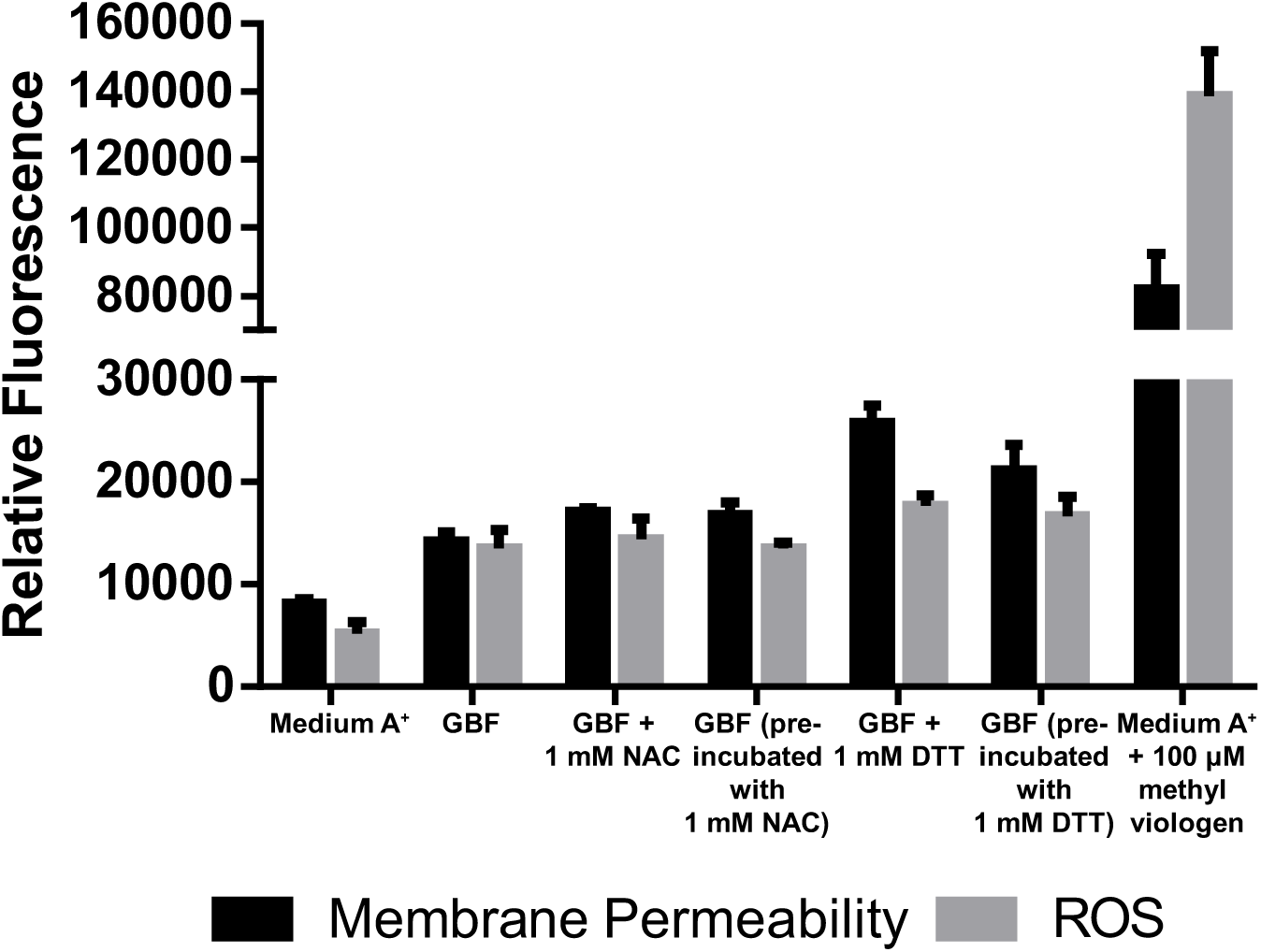
Reactive oxygen species and membrane permeability assay. Cells were grown to early linear phase in Medium A^+^ or 12.5% GBF with the appropriate treatment under continuous illumination (200 µmol photons m^−2^ s^−1^) with 1 % CO_2_ at 37°C. Fluorescence values were normalized to OD_730._ The values represent the mean ± SD of biological triplicates.

Photobleaching of the photosynthetic pigments is also a common symptom of oxidative stress in photosynthetic organisms and is caused by the accumulation of ROS (45). To investigate the effects of GBF media on the photosynthetic pigmentation, we performed whole cell absorbance scans of cultures cultivated at two different temperatures (27°C vs 37°C) and the same light intensity (200 µmol photons m^−2^ s^−1^). High GBF concentrations yielded enhanced chlorophyll, phycobilisome, and carotenoid degradation at 37°C (Figure 7a). At 27°C, photosynthetic pigments maintained intact relative to the control, regardless of GBF concentration, implying less ROS production at this temperature (Figure 7b).

**Figure 7.**
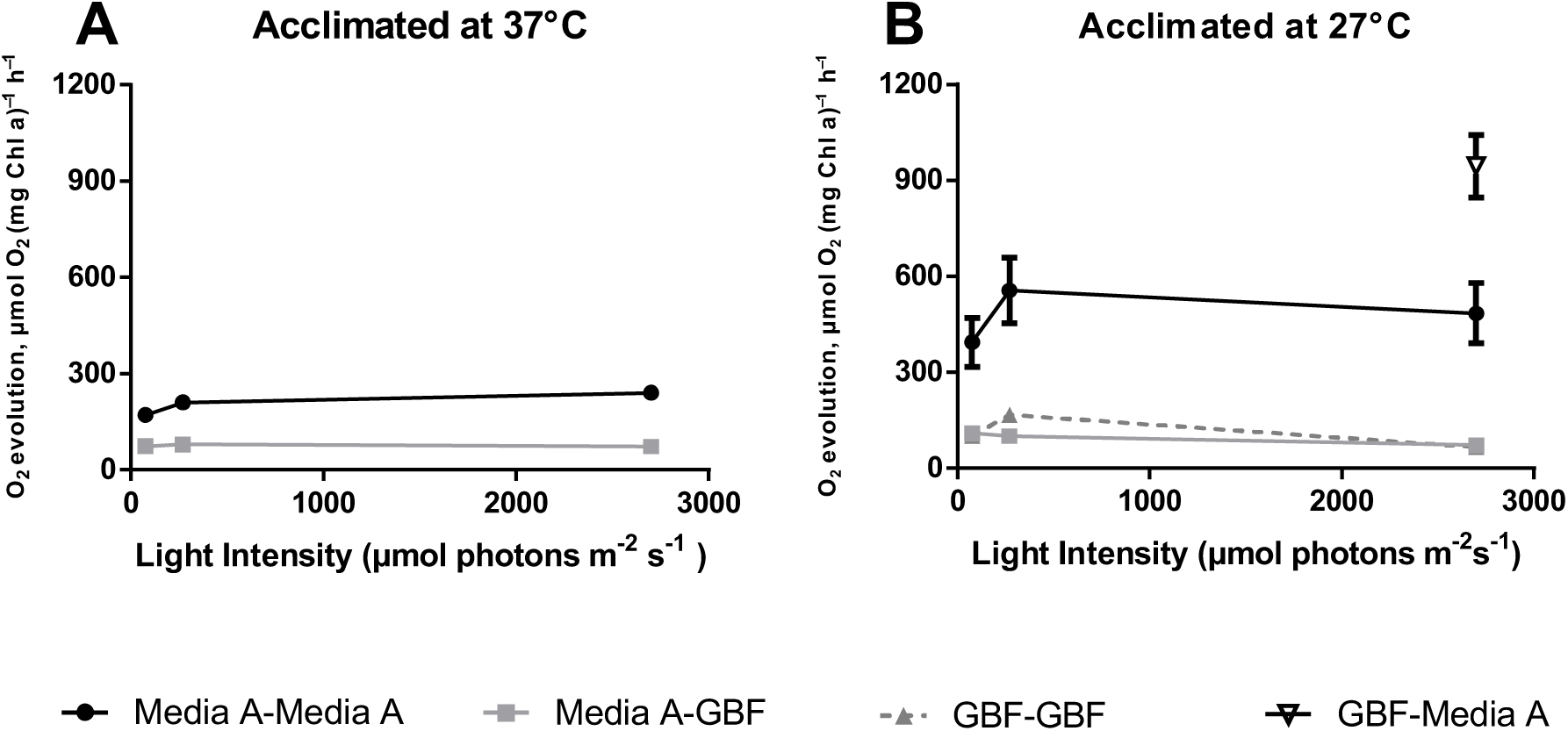
Rates of oxygen evolution as a function of acclimation temperature and light intensity. Cells were grown to early linear phase in Medium A^+^ or 12.5% GBF under continuous illumination (200 µmol photons m^−2^ s^−1^) with 1 % CO_2_ at (A) 37°C or (B) 27°C. Cells were pelleted, resuspended in Medium A^+^ or 12.5% GBF, and the rate of maximal oxygen evolution was measured with 10 mM HCO_3_^−^ as an electron acceptor at increasing light intensities. The values represent the mean ± SE of biological duplicates.

### Exposure to GBF retards oxygen evolution

To better delineate the cause of initial toxicity associated with high GBF concentrations, we measured maximal oxygen evolution rates for strains briefly exposed to media with 12.5% v/v GBF while increasing the light intensity. Measurements of photosynthetic oxygen evolution would allow for an indirect assessment of PSII activity and electron transfer (46). Cultures were grown to early linear phase in Medium A^+^ or GBF media, washed, and resuspended in the appropriate media. Resuspension media was saturated with HCO_3_^−^ (10 mM) in order to prevent inorganic carbon limitation. As expected, cells grown and assayed in Medium A^+^ at 37°C showed a clear increase in O_2_ evolution rate as the light intensity approached saturation at 2700 µm photons m^−2^ s^−1^, reaching a maximal rate of 240 ± 8 µmol O_2_ (mg Chl a)^−1^ h^−1^ (Figure 7a). Cells grown in Medium A^+^ at 37°C but assayed in 12.5% GBF had diminished O_2_ evolution rates at all light intensities, plateauing with a rate of 79 ± 6 µmol O_2_ (mg Chl a)^−1^ h^−1^ at an intensity of 270 µmol photons m^−2^ s^−1^ (Figure 7a). Thus, exposure to GBF under these conditions caused an immediate decrease in O_2_ production.

Next, we examined the effect of temperature in a similar experiment. Assays carried out in Medium A^+^ after growth in Medium A^+^ at 27°C showed much higher maximal O_2_ evolution rates at all tested light intensities than with cells grown at 37°C (Figure 7b), peaking at a rate 556 ± 102 µmol O_2_ (mg Chl a)^−1^ h^−1^). This was expected because elevated O_2_ evolution rates in low temperature grown cells have been previously reported and were attributed to a substantial change in photosystem stoichiometry (47). The O_2_ evolution rates of cells grown in Medium A^+^ at 27°C and then resuspended in 12.5% GBF stayed relatively constant at all of the tested intensities and were roughly 5-fold lower than in controls, with a maximal rate of 109 ± 22 µmol O_2_ (mg Chl a)^−1^ h^−1^ at an intensity of 75 µmol photons m^−2^ s^−1^. We compared the above rates to those from cultures grown in 12.5% GBF at 27°C to test if adaptation to GBF was met with changes in photosynthetic activity. At the examined light intensities, O_2_ evolution rates with 27°C GBF-adapted cultures were not statistically different from those measured in cultures grown at the same temperature in Medium A^+^. Finally, we conducted the inverse experiment, using cultures grown in GBF media at 27°C but assayed in Medium A^+^ under saturating light. Interestingly, they displayed the highest evolution rate of any tested condition, at 944 ± 96 µmol O_2_ (mg Chl a)^−1^ h^−1^ (Figure 7b). This suggested that there is a period of dynamic photosynthetic adaptation to overcome the stress of GBF, and that when the stress is removed, the cells have an enhanced capacity for photosynthetic activity.

Additional assays were performed at light saturation (2700 µmol m^−2^ s^−1^) with cultures adapted to 37°C (Figure 8) to further examine the effects of media conditions on oxygen evolution rates. Again, oxygen evolution rates in GBF (73 ± 8 µmol O_2_ (mg Chl a)^−1^ h^−1^) were much lower than Medium A^+^ (248 ± 21 µmol O_2_ (mg Chl a)^−1^ h^−1^). In order to rule out the effect of differences in overall ionic strength, we increased the osmolarity of the GBF medium by adding NaCl to match the levels found in Medium A^+^, and found no statistical difference in oxygen evolution as compared to experiments without added NaCl. Attempts to eliminate the inhibitory effect of GBF on oxygen evolution by gravity filtration through powdered activated carbon were unsuccessful. We also measured oxygen evolution rates in control experiments designed to inhibit photosynthetic electron transfer through 10 µM DCMU addition to Medium A^+^(48), and to prevent de-novo protein synthesis by pre-treating cultures with 800 µg ml^−1^ Lincomycin in Medium A^+^ (49), prior to incubation at light saturation for 1 hour. However, evolution rates with these controls were higher than those treated with GBF. Therefore, no definitive conclusion about the molecular mechanism responsible for reduced oxygen evolution rates in the presence of GBF could be made.

**Figure 8.**
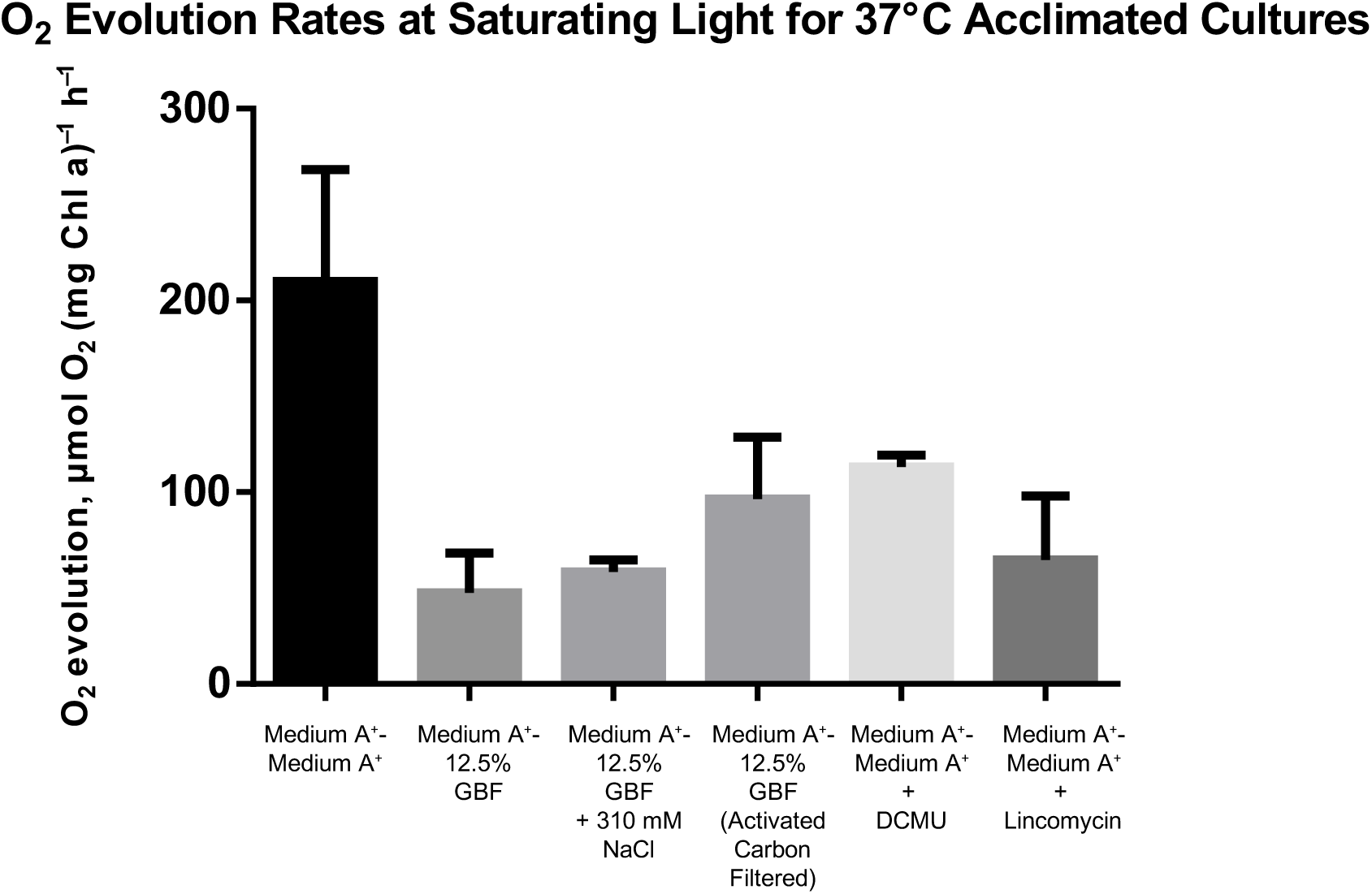
Photosynthetic oxygen evolution rates at saturating light conditions. Cells were grown to early linear phase in Medium A^+^ under continuous illumination (200 µmol photons m^−2^ s^−1^) with 1 % CO_2_ at 37°C). Cells were pelleted and resuspended in Medium A^+^, 12.5% GBF, 12.5% GBF + 310 mM NaCl (control for osmolarity), 12.5% GBF prefiltered with activated carbon, Medium A^+^ + 10 µM DCMU (control for inhibition for electron transport), or pre-treated with 800 µg ml^−1^ Lincomycin for 1 hour (control for the inhibition of protein synthesis.) Maximal oxygen evolution rates were measured with 10 mM HCO_3_ as an electron acceptor at 2700 µmol photons m^−2^ s^−1^. The values represent the mean ± SE of biological duplicates.

### Acclimation to GBF changes lipid content and composition

Based on the results described above, we hypothesized that the temperature dependent adaptation may be related to changes in membrane content and composition. We extracted total fatty acids of cultures grown at 27°C for 72 hours in 12.5% GBF or Medium A^+^, and analyzed the content after derivatization. We could detect and resolve all major saturated and unsaturated fatty acid species (Table 4). Cultures grown in GBF had greater totals of assayed fatty acid species (27 ± 7 mg FAME gDCW^−1^) when compared to cells grown in Medium A^+^ (15 ± 1 mg FAME gDCW^−^1) (Table 4). The most abundant fatty acid in all samples was 16:0 (42–45% of the total fatty acids). C18:2 ∆9,12 fatty acids comprised a significant fraction of Medium A^+^ grown cultures with 22% of the total fatty acid species. However, in GBF-grown cells C18:2 ∆9,12 fatty acids were only 15% of the total fatty acid pool, while C18:3 ∆9,12,15 fatty acids were twice as high in GBF grown cells (16%) than in Medium A^+^ grown cells (9%) (Table 4). These results suggest that cells were altering their membrane homeostasis when grown in GBF, as compared to standard growth in Medium A^+^. While we did not differentiate the fatty acid content and composition between the outer, plasma, or thylakoid membranes in our study, prior work done with the cyanobacterium *Synechocystis* PCC 6803 found similar fatty acid composition between thylakoid and plasma membranes (50).

**Table 4.** Fatty acid content and composition of cultures grown in Medium A^+^ or 12.5% GBF

media under continuous illumination (200 $\mu\text{mol photons m}^{-2} \text{s}^{-1}$ ) at 27°C for 72 hours
| Growth Conditions<sup>1</sup> |  | Fatty Acids Species |  |  |  |  |  |
| --- | --- | --- | --- | --- | --- | --- | --- |
|  |  | C16:0 | C16:1 Δ9 | C18:0 | C18:1 Δ9 | C18:2 Δ9,12 | C18:3 Δ9,12,15 |
| Medium A<sup>+</sup> | mg FA gDCW<sup>-1</sup> | 6.7 ± 0.7 | 2.0 ± 0.1 | 0.2 ± 0.1 | 1.5 ± 0.2 | 3.3 ± 0.2 | 1.4 ± 0.1 |
|  | % FA | 44 ± 1 | 13 ± 1 | 2 ± 0 | 10 ± 0 | 22 ± 1 | 9 ± 1 |
| GBF | mg FA gDCW<sup>-1</sup> | 11.6 ± 3.1 | 3.5 ± 0.8 | 1.3 ± 0.3 | 2.2 ± 0.7 | 3.9 ± 0.8 | 4.4 ± 1.6 |
|  | % FA | 43 ± 0 | 13 ± 0 | 5 ± 0* | 8 ± 0 | 15 ± 1* | 16 ± 1* |

<sup>1</sup> Strains were grown in the Medium A<sup>+</sup> or 12.5% GBF media under continuous illumination (200 $\mu\text{mol photons m}^{-2} \text{s}^{-1}$ ) at 27°C for 72 hours. Fatty acids were extracted and derivatized as previously described. The values represent the mean ± SD of two independent experiments. \* represent statistically significant differences from control (p-value<0.005 using an unpaired t-test)

## DISCUSSION

The amount and distribution of arable land and potable water are projected to change over the next several decades due to climate change, while the rise in global population and standard of living are expected to increase demand for these resources (51). Microalgal cultivation integrated with industrial and municipal wastewater treatment circumvents many of the resource concerns raised over biofuel production (2), while simultaneously removing additional nutrients and pollutants present in the wastewater (52). *Synechococcus* PCC 7002 is an intriguing possible platform for this purpose, since it can be readily engineered to produce high-value chemicals (53). However, under standard environmental conditions we were unable to obtain robust growth of PCC 7002 using a diluted municipal side stream (GBF) as a nutrient source. We hypothesized that this effect may be due to the presence of DOM, which has been demonstrated to cause a decrease in photosynthetic performance in various cyanobacteria strains (54–56). We investigated the effects of light intensity and temperature on the physiology of cultures grown in GBF, in an effort to find conditions conducive to high growth rates and biomass generation, and to better understand the mechanisms of GBF toxicity.

We propose that the herbicidal effect of GBF is primarily due to PSII inhibition as shown in Figures 7 and 8, brought about by quinone or phenolic compounds within the “soluble microbial products” found in our EEM scans (Figure 2). Humic material enriched in quinone and phenolic compounds can inhibit cyanobacterial growth (57, 58). Phenolic photosynthetic electron transfer inhibitors, such as 2,5-Dibromo-3-methyl-6-isopropyl-p-benzoquinone (DBMIB), alter the redox potential of the PQ pool of PSII by blocking forward electron transfer to the cytochrome b_6_/f complex (59). The immediate decrease in oxygen evolution (Figure 7) and high rates of initial membrane permeability (Figure 4) upon GBF exposure suggests that the toxic compound(s) rapidly cross the outer membrane, where it subsequently interrupts photosynthetic electron flow. Chronic exposure to phenolic herbicides eventually leads to radical-catalyzed back reactions that trigger ROS formation (60) (Figure 5) that facilitates complex destruction (Figure 6) and cell death (45, 61–64). Some phenolic herbicides may also act as arylating agents, causing covalent binding to macromolecules via Michael addition and thiol pool depletion (65). The inability of the reducing agents we tested to maintain membrane integrity suggests that GBF-induced cytomembrane permeability is likely caused by redox cycling, and not arylation.

**Figure 6.**
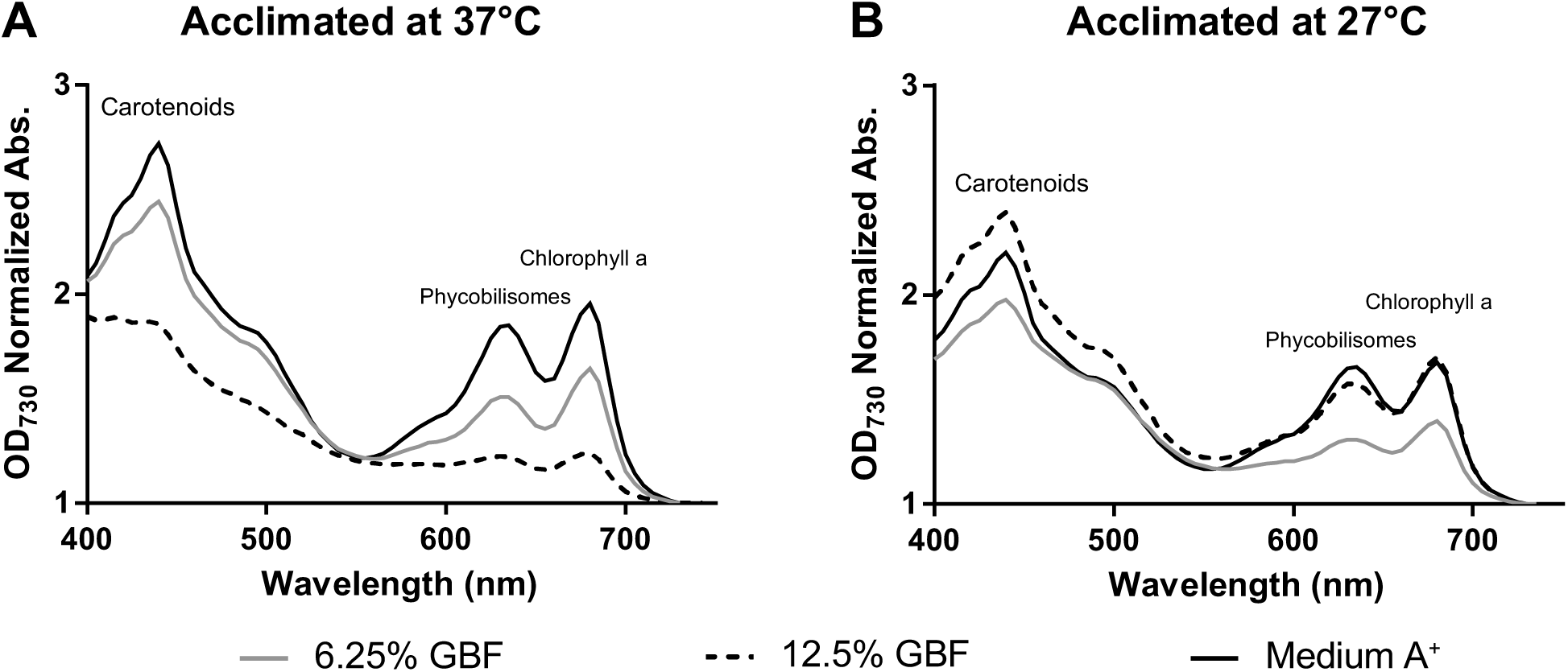
Absorption spectra of PCC 7002 cultures exposed to GBF, at two different cultivation temperatures. Cultures were grown in Medium A^+^, 6.25 %, or 12.5 % (v/v) GBF media under continuous illumination (200 µmol photons m^−2^ s^−1^) with 1 % CO_2_ at (A) 37°C or (B) 27°C for 72 hours. The spectra were recorded in dilute cell suspensions and normalized to an OD_730_. The peak at 438 nm is due to carotenoids, the peak at 637 nm is due to phycobilisomes, and the peak at 683 nm is due to chlorophyll a.

We found that cultivation temperature was an important factor determined whether PCC 7002 could grow under high GBF concentrations. Numerous physiological processes are altered at low temperatures (66). Notably, the fatty acyl chains in membranes undergo a transition from a fluid to nonfluid state (67). Cyanobacterial membrane organization is complicated by the simultaneous existence of the outer, plasma, and thylakoid membranes, each with a designated physiological function (68). As part of the homeoviscous response upon a shift to a lower temperature, cyanobacteria alter the expression of desaturases in their plasma and thylakoid membranes (69) to increase unsaturated fatty acid content and maintain optimal membrane function (70). Optimal thylakoid membrane fluidity is a critical factor in photosynthetic electron transport, because co-utilized redox components must be able to move quickly between photosynthetic and respiratory complexes to ensure ideal electron flow (71, 72). Temperature is known to influence the kinetics governing the redox state of plastiquinone (PQ) through alterations in thylakoid membrane composition and fluidity (73). We hypothesize that alteration of the thylakoid membrane to circumvent GBF induced changes in the overall redox state is also an important component of the adaptive response to lower temperatures, given the increase in total unsaturated fatty acids observed during cultivation in GBF-based media (Table 4). Upon shifts to a lower temperature (74), PCC 7002 has also been shown to translocate the phycobilisomes from PSII to PSI, thereby decreasing the ratio of reducing power to proton-motive force (75). Mutant cells of the cyanobacterium *Synechocystis* PCC 6803 that lacked polyunsaturated fatty acids were unable to perform these state transitions at low temperatures (76). This change in photosystem arrangement at 27°C may also contribute to acclimation to the photoinhibitory compound. Future efforts to increase tolerance to these toxicants might include disruption of the inhibitor binding sites (77, 78) or optimizing thylakoid membrane fluidity (79).

## CONCLUSIONS

Under standard cultivation conditions of 37°C with the cyanobacteria *Synechococcus* sp. strain PCC 7002, there was a dose-dependent relationship between liquid anaerobic digestate (GBF) concentration and membrane permeability. This dose-dependent relationship mirrored ROS production and photopigment degradation. Digestate contained dissolved organic matter constituents that were likely affecting photosynthetic electron transport. Decreasing the cultivation temperature to 27°C enabled robust cultivation at high digestate concentrations, resulting in high biomass productivities. This temperature dependent tolerance may be due to changes in membrane properties. Our study highlights an unanticipated influence of dissolved organic matter on photosynthetic growth and physiology in wastewater based media, as well as a potential mechanism for tolerance in low temperatures.

## ACKNOWLEDGEMENTS

This work was funded by the US National Science Foundation (EFRI-1240268). TCK is the recipient of a National Institutes of Health (NIH) Biotechnology Training Fellowship (NIGMS-5 T32 GM08349) and a fellowship from the UW-Madison College of Engineering's Graduate Engineering Research Scholars (GERS) program. The authors are grateful to Richard Mikel, Matthew Dysthe, and Derek Jacobs for help with routine sampling, and Andrew Maizel and Christina Remucal for advice on the excitation−emission matrix (EEM) fluorescence spectroscopy.

